# The Suprachiasmatic Nucleus Regulates Anxiety-Like Behavior in Mice

**DOI:** 10.1101/2020.05.08.085217

**Authors:** Chelsea A. Vadnie, Lauren A. Eberhardt, Mariah A. Hildebrand, Hui Zhang, Darius Becker-Krail, Lauren M. DePoy, Ryan W. Logan, Colleen A. McClung

## Abstract

Circadian rhythms are commonly disrupted in individuals with depression and/or anxiety disorders. Animal studies indicate that circadian rhythm disruption can cause increased depressive and anxiety-like behavior, but the underlying mechanisms are unclear. Currently, there is conflicting evidence as to whether the master pacemaker in the brain, the suprachiasmatic nucleus (SCN), plays a key role in regulating psychiatric-related behavior. To investigate the role of the SCN in regulating depressive and anxiety-like behavior in mice, we directly manipulated the neural activity of the SCN using two chronic optogenetic stimulation paradigms. Repeated stimulation of the SCN late in the active phase (circadian time 21, CT21) shortened the period and dampened the amplitude of homecage activity rhythms. Repeated stimulation of the SCN at unpredictable times during the dark phase dampened, fragmented and reduced the stability of homecage activity rhythms. In both SCN optogenetic stimulation paradigms, dampened homecage activity rhythms (decreased amplitude) was associated with increased measures of anxiety-like behavior, but not in control mice. Increased fragmentation and decreased day-to-day stability of homecage activity also correlated with increased anxiety-like behavior. Unexpectedly the change in period of homecage activity rhythms was not directly associated with any psychiatric-related behavior. Furthermore, we did not observe consistent correlations between homecage activity amplitude and depressive-like behavior in stimulated mice. Taken together, these results indicate that SCN-mediated dampening of rhythms is directly correlated with increased anxiety-like, but not depressive-like behavior in mice. This work is an important step in understanding how specific SCN neural activity disruptions affect mood and anxiety-related behavior.

## Introduction

Circadian rhythms are physiological processes that oscillate with an approximate 24-hour period. Circadian rhythms are endogenously generated and can entrain to environmental cues, such as changes in the light/dark (LD) cycle. At the cellular level, rhythms are generated by transcriptional/translational feedback loops of circadian genes, which are expressed in nearly every cell in the body^1^. The suprachiasmatic nucleus (SCN) in the hypothalamus acts as the master pacemaker, synchronizing bodily rhythms with each other and the environment. The SCN is an effective pacemaker because neurons in the SCN are highly coupled and exhibit a robust circadian rhythm of neuronal firing^2^. Specifically, SCN neuronal firing is high during the subjective day and low during the subjective night in both nocturnal and diurnal species^3,4^. Additionally, the SCN receives light information from intrinsically photosensitive retinal ganglion cells (ipRGCs) via glutamatergic projections^5,6^. Thus, the pattern of SCN neuronal firing can be altered by both environmental and genetic insults, ultimately disrupting circadian rhythms^7-9^. Disrupted circadian rhythms are associated with adverse health outcomes, but it is unclear whether the SCN specifically plays a role in regulating mental and physical health.

Circadian rhythms are frequently disrupted in psychiatric disorders where rhythms are typically dampened or shifted^10-13^. For some individuals, environmental circadian disruption, such as shift work and jet lag, can trigger or exacerbate symptoms of psychiatric disorders^14-16^. Genetic variations in the molecular clock are also associated with psychiatric disorders, such as bipolar disorder, anxiety and depression^17-19^. Furthermore, treatments that directly affect rhythms, such as social rhythm and bright light therapy, have therapeutic effects for some individuals with psychiatric disorders^20,21^. Taken together, this suggests that circadian rhythm disruption plays a causative role in some individuals with depression and/or anxiety, but the underlying mechanisms are unclear.

There is conflicting evidence as to whether the SCN directly modulates mood and anxiety. Several human postmortem brain studies find alterations in specific neurotransmitters and/or receptors in the SCN of individuals with a mood disorder^22-24^. Human postmortem work has also shown that gene expression rhythms are weaker and desynchronized in brain regions outside of the SCN in major depressive disorder^25^. Similarly, in mice, we have found that disrupting circadian gene expression in brain regions outside of the SCN can increase depressive and anxiety-like behavior^26,27^. One possibility for how this might occur naturally is that environmental disturbances could disrupt SCN rhythms which in turn disrupt gene expression rhythms throughout the brain, potentially increasing depressive and anxiety-like behavior. In animal models, there is evidence to either support or refute that the SCN regulates depressive and/or anxiety-like behavior. Knocking down *Bmal1* expression in the SCN was shown to increase depressive and anxiety-like behavior in mice^28^. However, recent work shows that light through an ipRGC-brain pathway independent from the SCN increases depressive-like behavior in mice, suggesting that the SCN does not play a major role in regulating light-induced depressive-like behavior^29^. Contrary to both of these findings, the behavioral effects of lesioning the SCN generally leads to an antidepressant response, i.e. less immobility in the forced swim test (FST)^30,31^. One reason for these conflicting results may be that not one of the above studies directly manipulated the neural activity of the SCN.

It is clear from animal studies that decreasing the amplitude or chronically shifting rhythms can increase anxiety and/or depressive-like behavior. For example, repeated advances in the LD schedule, constant light and non-24 hr LD cycles can all disrupt rhythms and increase depressive and/or anxiety-like behavior in rodents^32-37^. Previously, we found that unpredictable chronic mild stress, a well-established paradigm to increase depressive and anxiety-like behavior, dampened locomotor activity, body temperature, and SCN rhythms in mice^38^. Interestingly, dampened SCN PERIOD2::LUCIFERASE (PER2::LUC) rhythms were directly correlated with increased depressive and anxiety-like behavior, suggesting a role for dampened SCN rhythms in mediating the behavioral effects of stress. However, since light and stress manipulations can affect the brain in SCN-independent ways there is a need to understand whether SCN-mediated shifting and/or dampening of rhythms increases depressive and anxiety-like behaviors. If we find that specific SCN manipulations can cause psychiatric-related behaviors, this could lead to the development of new therapeutics for anxiety or mood disorders that directly target the SCN.

To determine how SCN-mediated shifting and dampening of rhythms affects psychiatric-related behaviors in mice, we used optogenetics to increase SCN neuronal firing at specific times of day. We utilized a previously described SCN optogenetic stimulation paradigm^39^ which was shown to increase SCN neuronal firing and phase shift SCN PER2::LUC rhythms similar to what would be expected for the effects of environmental light on the SCN^39^. Here, our goal was to use a similar SCN optogenetic stimulation paradigm to repeatedly shift or dampen rhythms to determine the specific role of the SCN in regulating depressive and anxiety-like behaviors.

## Materials and methods

### Animals and housing conditions

Homozygous *Vgat*-Cre mice (B6.FVB.129S6-*Slc32a1*^*tm2(cre)Lowl*^/J; The Jackson Laboratory; stock 016962) were crossed with homozygous Cre-dependent ChR2 mice (Ai32, B6.129S-*Gt(ROSA)*^*26Sortm32(CAG-COP4*H134R/EYFP)Hze*^/J; The Jackson Laboratory; stock 012569) to yield heterozygous *Vgat*;ChR2 mice. We used male heterozygous *Vgat*;ChR2 and homozygous *Vgat*-Cre mice (8-15 wks old) for experiments. Mice were maintained on a 12:12 LD schedule (lights on at 7 AM and off at 7 PM, except where described) and were provided with food and water *ad libitum*. All animal use was conducted in accordance with the National Institute of Health guidelines and approved by the Institutional Animal Care and Use Committees of the University of Pittsburgh. Sample sizes were chosen based on adequately powered sizes in our previous studies.

### Surgery

Mice were anesthetized with isoflurane and placed in a stereotaxic device. An optic fiber (NA 0.39, 400 µm core, Thorlabs, Newton, NJ) coupled to a metal ferrule was implanted proximally dorsal to the SCN (AP −0.1 mm, ML 0.0 mm, DV −5.0 mm). Optic fiber implants were adhered to the skull with cement (C&B Metabond, Parkell, Edgewood, NY; black dental cement, Lang Dental Manufacturing, Wheeling, IL). The incision was closed with tissue adhesive (Vetbond, 3M, St. Paul, MN). Light transmission was measured before implantation and postmortem. Only mice with fibers with ≥ 80% efficiency were used.

### Circadian activity and sleep/wake recordings

Homecage activity and sleep/wake measurements were determined by piezoelectric recording of movements and breath rate (PiezoSleep, Signal Solutions LLC, Lexington, KY). Mice were individually housed in four-cage unit polycarbonate boxes in a ventilated, light-controlled cabinet. Mice were randomly assigned to experimental groups to counterbalance cabinet and box position. Each cage rested on a polyvinylidene difluoride (PVDF) square sensor (17.8 × 17.8 cm, 110 µm thick) protected by a thin plastic tray (50.8 µm)^40^. A rubber pad between each sensor and the base prevented crosstalk between the cages. The sensors were connected to an amplifier. Pressure signals and breath rates were classified as movements related to activity and inactivity or sleep and wake.

### *In vivo* light delivery

A 100 mW 473 nm DPSS laser and a 100 mW 447 nm diode laser (OEM Laser Systems, Midvale, UT) were connected to commutators (Doric Lenses Inc., Quebec, Canada) connected to multi-mode fiber optic black-jacketed patch cords (NA 0.22, 200 µm core, Doric Lenses Inc.). Mice were removed from their home cages using a dim red head lamp. Mice were attached to a patch cord and placed in black polycarbonate 20 cm^3^ box. Heat shrink tubing blocked blue light leakage from the connection between the patch cord and fiber implant. For all experiments, mice received 1 hr sessions of blue light pulses (8 Hz, 10 ms pulse width, light intensity at the fiber tip 8-11 mW) or as a control were sham stimulated (connected to patch cords, but did not receive blue light pulses). Light spread and power densities (mW/mm^2^) were calculated as described previously^41^. Mice that lost fiber implants, had poor fiber placement, or had low fiber efficiency were removed from the study.

### Chronic SCN optogenetic stimulation at CT21

*Vgat*;ChR2 mice were habituated to a reverse LD schedule (lights off at 5 AM and on at 5 PM) for 16 days and were then placed in constant darkness (DD) for 5 days to measure baseline free-running rhythms. To shift rhythms, mice received SCN stimulation at circadian time 21 (CT21) every 3 days, where CT12 was defined as activity onset. Based on environmental light studies^42,43^, we expected direct optogenetic stimulation of the SCN at CT21 to increase SCN neuronal firing and advance rhythms. For the first stimulation day, stimulation time was determined by finding the onsets of activity on DD days and using linear regression (ClockLab, Actimetrics, Wilmette, IL) to predict the onset for the first stimulation day. For subsequent days, stimulation times were determined by finding the onsets for the 2 days in between stimulations. Behavior testing took place every 3 days during the active phase, following the 6^th^ stimulation as described below. Stimulations continued every 3 days on non-behavior testing days.

To ensure that blue light pulses in the absence of ChR2 was not capable of altering homecage activity rhythms, we implanted optic fibers in a separate group of *Vgat*;ChR2 as well as control, *Vgat*-Cre mice. The experiment was conducted as described above except mice only received 6 total SCN stimulations.

### Chronic SCN optogenetic stimulation at unpredictable times during the dark phase

*Vgat*;ChR2 mice were habituated to a reverse LD schedule (lights off at 5 AM and on at 5 PM) for 10 days before baseline rhythms were measured for 7 days. To dampen the amplitude of rhythms, mice received daily SCN optogenetic stimulation at unpredictable times during the dark phase for 8 days. Since SCN neuronal activity is low during the dark phase, stimulations should increase the trough of SCN neuronal activity rhythms and subsequently dampen the amplitude of rhythms. We stimulated at unpredictable times to prevent entrainment to the stimulations. Behavior testing took place on the 9^th^ day during the inactive phase as described below. Daily SCN stimulations continued at unpredictable times during the dark phase throughout behavior testing. The amplitude of activity rhythms was monitored to ensure efficacy of optogenetic stimulations and mice which showed a significant amplitude change moved forward into additional behavioral studies.

### Behavioral Assays

For stimulations at CT21, behavior testing occurred between CT14-18, during the active phase. Mice were given 30 min to habituate to the room and all behavior testing took place under dim red light (< 10 lux).

For unpredictable dark phase stimulations, behavior testing took place during the light phase (ZT3-6). Mice were given 1 hr to habituate to the room before testing. Behavior testing for the open field (OF) and elevated plus maze (EPM) took place under dim white light (∼20 lux) to promote exploratory behavior. Other behavior tests occurred under standard room lighting.

### Statistical analysis

We quantified homecage activity using WakeActive and ActivityStatistics software, and sleep using SleepStats2p10 software (Signal Solutions LLC). For homecage activity analysis, data were visualized as normalized actograms with 3 min bin sizes in ClockLab. The period of activity rhythms was determined by applying a least-squares fit to the onset measures. Homecage activity amplitude was quantified using a least-squares fit cosinor analysis. For stimulation at CT21, the period of homecage activity over the analysis range was used for the cosinor fit. For unpredictable dark phase stimulation, a tau of 24 hrs was used. To obtain additional measures of activity disruption after unpredictable dark phase stimulation, we carried out a non-parametric circadian rhythm analysis in ClockLab to measure relative amplitude, daily fragmentation (intradaily variability) and day-to-day stability (interdaily stability).

The change in homecage activity and sleep measures relative to baseline were determined for each animal. The behavior testing period was not included in the homecage activity and sleep analysis since the mice were removed from the boxes for extended periods of time. Unpaired two-tailed *t*-tests were used to assess differences in measures between the control and stimulated *Vgat*-Cre;ChR2 mice. Where there was unequal variance Welch’s *t*-tests were used and where there was not a normal distribution Mann-Whitney tests were used. Two-way ANOVAs followed by Tukey post-hoc tests where appropriate were used to determine differences in measures for the control experiment using *Vgat*-Cre and *Vgat*-Cre;ChR2 mice. Correlations between homecage activity measures and behavior were assessed using two-tailed Pearson correlations. Data were analyzed using Prism 7 for PC (GraphPad Software, San Diego, CA). Bar graphs are presented as mean ± SEM and *p* < 0.05 was considered significant.

See Supplementary Methods for additional details.

## Results

### Chronic SCN optogenetic stimulation at CT21 shortens the period and dampens the amplitude of homecage activity rhythms

To selectively stimulate neurons in the SCN, we generated heterozygous *Vgat*;ChR2 mice. VGAT is a vesicular transporter that loads GABA and glycine into synaptic vesicles. Since nearly all SCN neurons are GABAergic^44^, this was an effective approach to obtain high ChR2 expression in the SCN (**Supplementary Fig. S1a**). Optic fibers were implanted proximally dorsal to the SCN. High expression of ChR2 in the SCN coupled with limited spread of blue light in the brain allowed us to specifically target the SCN (**Supplementary Fig. S1b**).

To determine whether optogenetic stimulation of the SCN increases SCN neuronal activity, we quantified the number of c-Fos-positive cells in the SCN of *Vgat*;ChR2 mice after stimulation or sham stimulation. Consistent with previous work showing that a similar SCN optogenetic paradigm increased SCN neuronal firing^39^, SCN stimulation at CT21 increased the number of c-Fos-positive cells in the SCN relative to controls (*t*_(9)_=7.91, *p*<0.0001, **Fig. 1a-b**).

**Figure 1.**
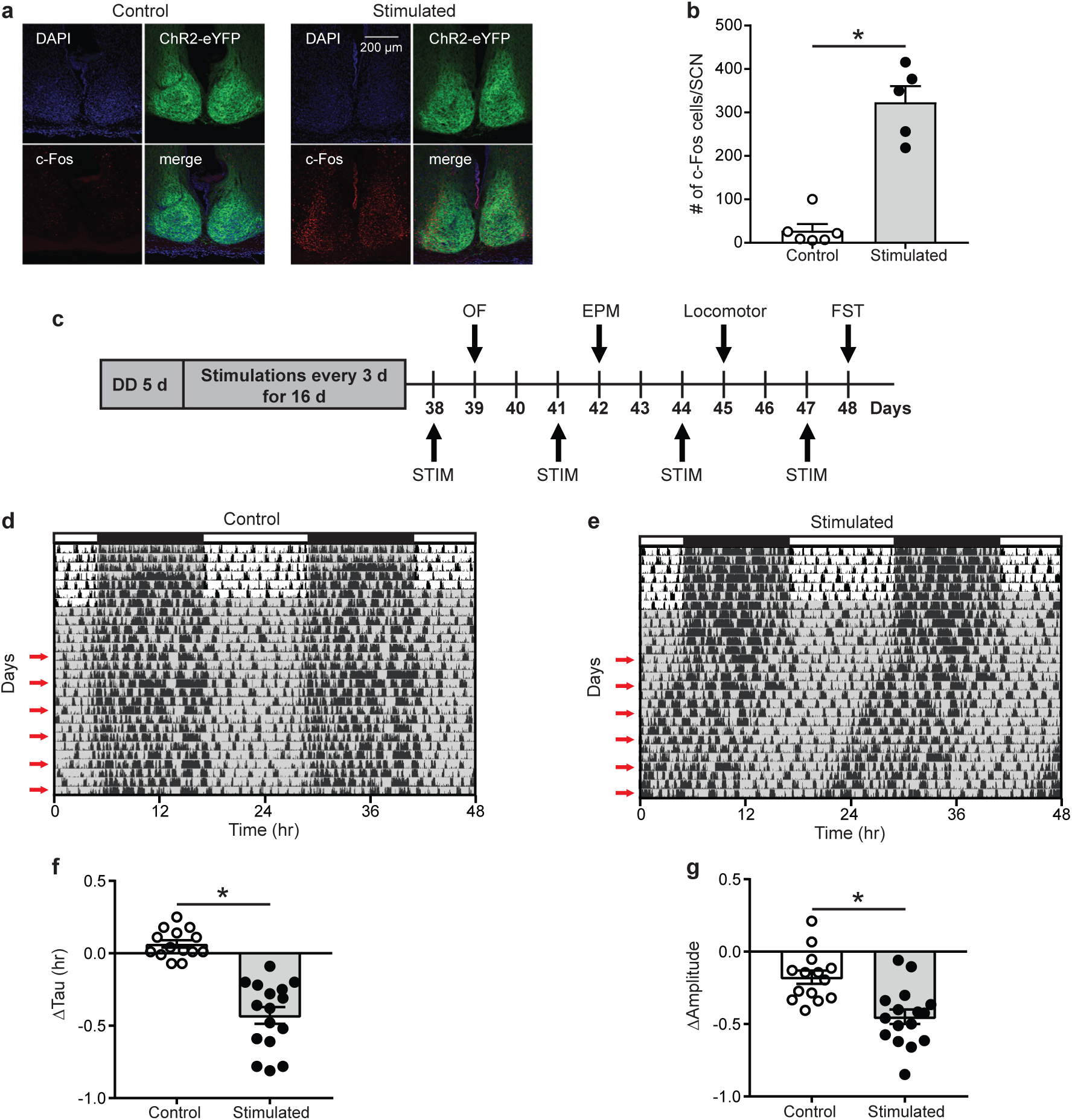
Effects of SCN stimulation at CT21 on c-Fos cells in the SCN and homecage activity rhythms. (a) Representative confocal images of coronal mouse brain slices containing the SCN from a control and stimulated mouse. (b) The average number of c-Fos-positive cells per mouse SCN were increased after stimulation at CT21. *n* = 5-6 mice. (c) Experimental design to determine the effects of repeated SCN stimulation at CT21 on psychiatric-related behaviors. stimulation (STIM). (d-e) Representative double-plotted actograms of homecage activity of a control and stimulated mouse. Gray shading indicates when lights were off and red arrows indicate the days when stimulations or sham stimulation occurred at CT21. Change in homecage activity circadian parameters were measured relative to baseline in DD. (f) Stimulated mice exhibited a greater reduction in homecage activity period or tau. (g) Stimulated mice showed a greater decrease in homecage activity amplitude. \**p* < 0.05. *n* = 14-16.

Next, we wanted to determine whether chronic stimulation of the SCN at CT21 would result in advancing homecage activity rhythms (**Fig. 1c, Supplementary Fig. S1c**). Stimulation at CT21 shortened the homecage activity period or tau relative to controls (*t*_(20.76)_=7.79, *p*<0.0001, **Fig. 1d-f**). Consistent with studies showing that shifts in the LD schedule can dampen the amplitude of rhythms^45,46^, SCN stimulation at CT21 also dampened the amplitude of homecage activity rhythms relative to controls (*t*_(28)_=4.01, *p*<0.001, **Fig. 1g**). The slight reduction in amplitude of the control mice relative to baseline (−0.176 ± 0.046) was likely due to interruption in homecage recording for sham stimulation and handling stress. We did not observe a significant change in the average daily percent time mice spent sleeping (**Supplementary Fig. S2a**, *U*=0.96, *p*=0.52). However, there was a reduction in average daily sleep bout duration relative to controls (**Supplementary Fig. S2b**, *t*_(23.74)_=2.11, *p*<0.05). Together our data suggest that chronic SCN stimulation at CT21 altered the pattern of homecage activity with no effect on total sleep time.

As a control, we stimulated or sham stimulated *Vgat*-Cre or *Vgat*-Cre;ChR2 mice to ensure blue light pulses in the absence of ChR2 was not affecting homecage activity rhythms. Only mice that expressed ChR2 and received blue light pulses showed a significant shortening (genotype x stimulation, *F*_(1,24)_=39.68, *p*<0.0001) and dampening of homecage activity rhythms (genotype x stimulation, *F*_(1,24)_=4.94, *p*<0.05), confirming that we were not observing effects of blue light pulses alone on rhythms (**Supplementary Fig. S3**).

### Dampened amplitude of homecage activity rhythms after SCN stimulation at CT21 was associated with increased anxiety-like behavior in the OF

To determine whether SCN-mediated advancing of rhythms causes alterations in psychiatric-related behaviors, we examined correlations between homecage circadian activity measures and behavior. In stimulated mice only, dampened homecage activity rhythms relative to baseline were correlated with fewer entries and less time in the center of the OF (**Fig. 2a-b**). Distance traveled in a novel environment was not correlated with change in homecage activity amplitude (**Fig. 2c**), indicating that correlations with OF behavior were independent from any effect of SCN stimulation on general locomotion. In control mice, there were no correlations between change in homecage activity amplitude and behavior in the OF or locomotion in a novel environment (**Fig. 2d-f**). In stimulated mice, there was a strong trend for more dampened homecage activity rhythms to correlate with fewer percentages of open arm entries (**Fig. 2g**). However, a similar trend was observed in controls (**Fig. 2j**) and no correlations were observed with open arm time (**data not shown**). Surprisingly, we did not observe correlations between changes in the period of homecage activity rhythms and behavior (**Supplementary Fig. S4**). Taken together, dampened homecage activity rhythms due to chronic SCN stimulation at CT21 are associated with increased anxiety-like behavior in the OF.

**Figure 2.**
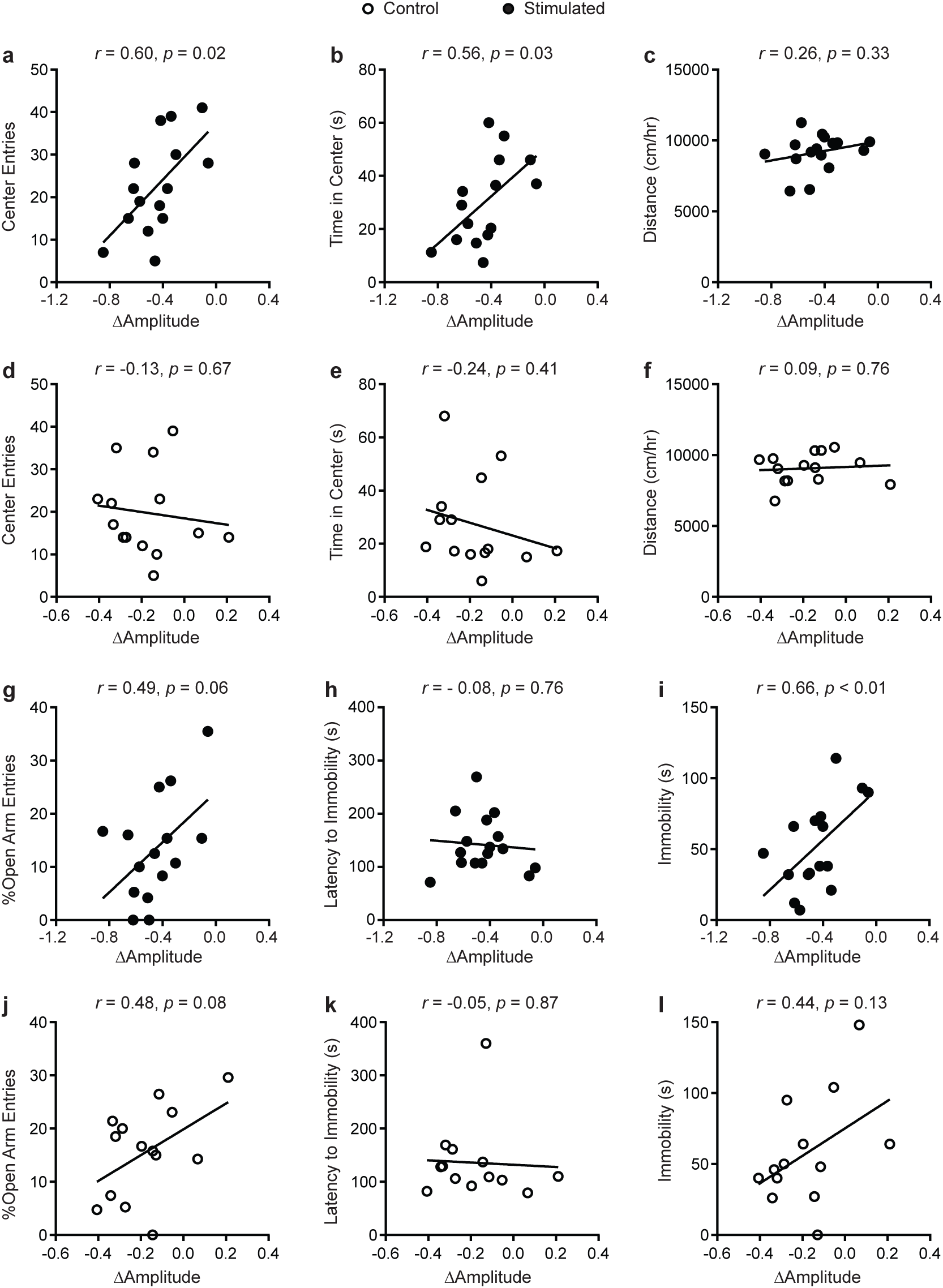
Correlations between change in homecage activity amplitude and behavior in mice that received stimulations or sham stimulations of the SCN at CT21. (a-b, d-e) OF, (c, f) locomotor, (g, j) EPM, (h-i, k-l) FST. A dampened amplitude of homecage activity rhythms in stimulated mice was correlated with increased anxiety-like behavior in the OF and decreased immobility in the FST. *n* = 13-16.

In stimulated mice (**Fig. 2h-i**), dampened homecage activity amplitude also correlated with reduced immobility in the FST but not latency to immobility. In control mice, change in homecage activity amplitude was not correlated with behavior in the FST (**Fig. 2k-l**). Together this suggests that SCN-mediated dampening of rhythms may reduce depressive-like behavior.

### Unpredictable stimulation of the SCN during the dark phase dampened the amplitude of homecage activity rhythms

To more specifically determine the effects of SCN-mediated disruption of the amplitude of rhythms on mood and anxiety-related behaviors, we kept mice on a 12:12 LD schedule and stimulated the SCN daily at unpredictable times during the dark phase (**Fig. 3a**). Stimulated mice exhibited a significantly greater decrease in homecage activity amplitude relative to controls (**Fig. 3b-d**, *t*_(19)_ = 2.21, *p* = 0.04). The decrease in homecage activity amplitude in stimulated mice cannot be attributed to an overall decrease in homecage activity since we did not find a significant difference in midline estimating statistic of rhythm (MESOR) between the groups (**Fig. 3e**, *t*_(19)_ = 1.29, *p*=0.21). In addition, there were no differences in the change in average daily percent sleep (**Supplementary Fig. S5a**, *U*=48, *p*=0.65) or average daily sleep bout duration (**Supplementary Fig. S5b**, *t*_(19)_=0.86, *p*=0.40), indicating that the reduction in homecage activity amplitude in stimulated mice is not due to a change in total sleep.

**Figure 3.**
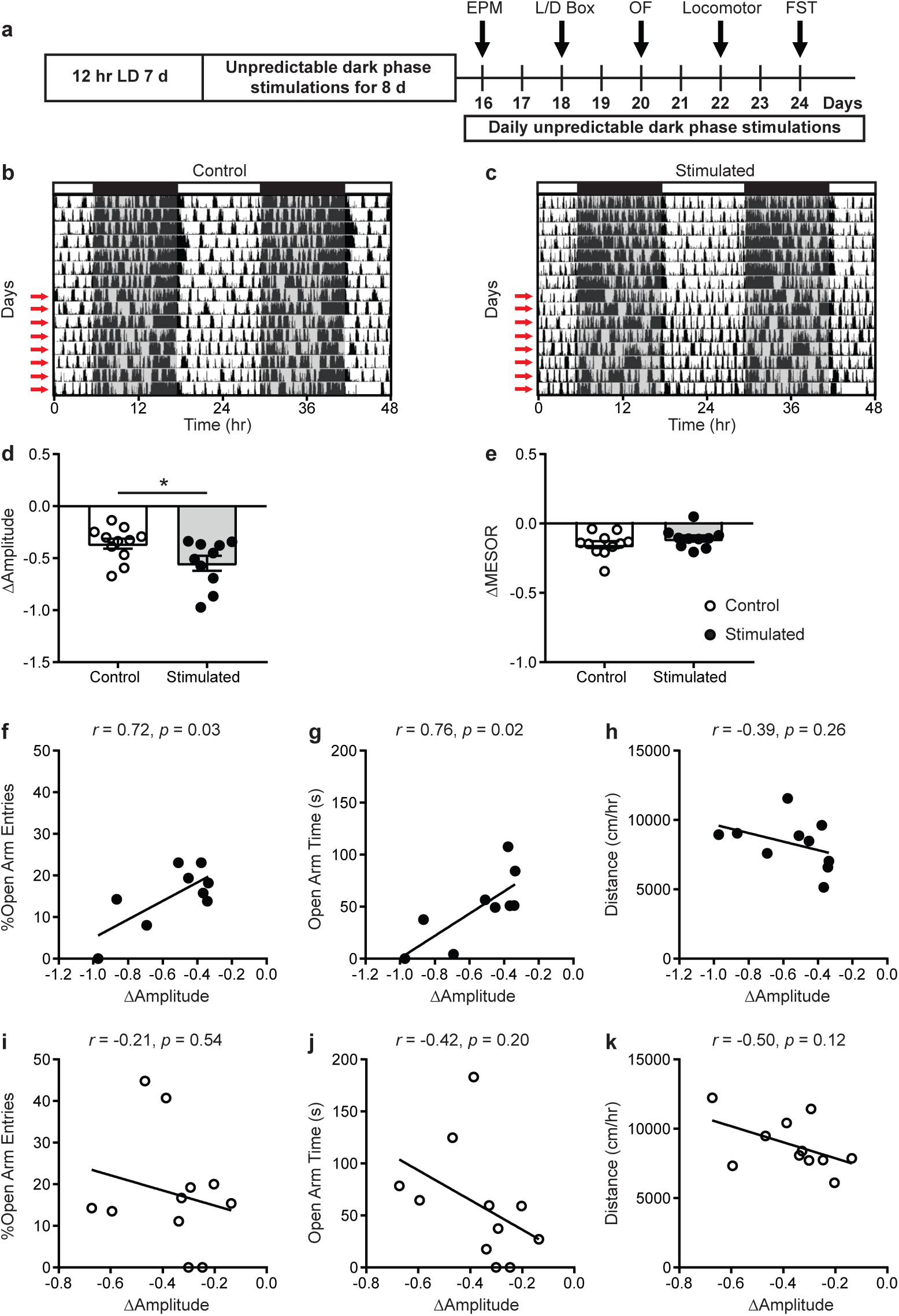
Homecage circadian activity changes and correlations with behavior in mice that received stimulations or sham stimulations of the SCN at unpredictable times during the dark phase. (a) Experimental design. (b-c) Representative double-plotted actograms of homecage activity of a control and stimulated mouse. Gray shading indicates when lights were off and red arrows indicate when the daily stimulations took place. Change in homecage activity circadian parameters were measured relative to baseline in LD. (d) Stimulated mice exhibited a greater reduction in homecage activity amplitude. (e) There was no difference in the change in midline estimating statistic of rhythm (MESOR) between control and stimulated mice. Correlations between change in homecage activity amplitude and (f-g, i-j) EPM and (h,k) locomotor behavior in mice that received daily unpredictable dark phase stimulations or sham stimulations of the SCN. \**p* < 0.05. *n* = 9-11.

### Dampened amplitude of homecage activity rhythms with unpredictable dark phase stimulation of the SCN was associated with increased anxiety-like behavior in the EPM

To determine how SCN-mediated dampening of rhythms affects anxiety and depressive-like behaviors, we performed correlations between the change in homecage activity amplitude and behavior. In stimulated mice, dampened homecage activity rhythms were strongly correlated with reduced open arm entries and open arm time in the EPM (**Fig. 3f-g**). Distance traveled in a novel environment was not correlated with the change in homecage activity amplitude in stimulated mice (**Fig. 3h**), indicating that the observed correlations were independent from any effect of SCN stimulation on general locomotion. In control mice, no correlations were observed between changes in homecage activity amplitude and behavior in the EPM or distance traveled in a novel environment (**Fig. 3i-k**). Surprisingly, in stimulated mice we did not observe correlations between homecage activity amplitude and other measures of anxiety-like behavior or behavior in the FST (**Fig. 4a-c, g-i**). Unexpectedly, in control mice, dampened amplitude of homecage activity rhythms was correlated with increased center entries and center time in the OF (**Fig. 4d-e**). There were also trends for correlations between dampened amplitude of homecage circadian rhythms and behavior in the LD box in control mice (**Fig. 4f&j**). Thus, in sham-stimulated mice, handling-induced dampening of homecage activity rhythms was associated with less anxiety-like behavior. In control mice, there were no correlations between change in the amplitude of homecage activity rhythms and behavior in the FST (**Fig. 4k-l**). Together our data suggest that dampening the amplitude of rhythms by unpredictable dark phase stimulation of the SCN increases anxiety-like behavior specifically in the EPM with no impact on depression-like behavior in the FST.

**Figure 4.**
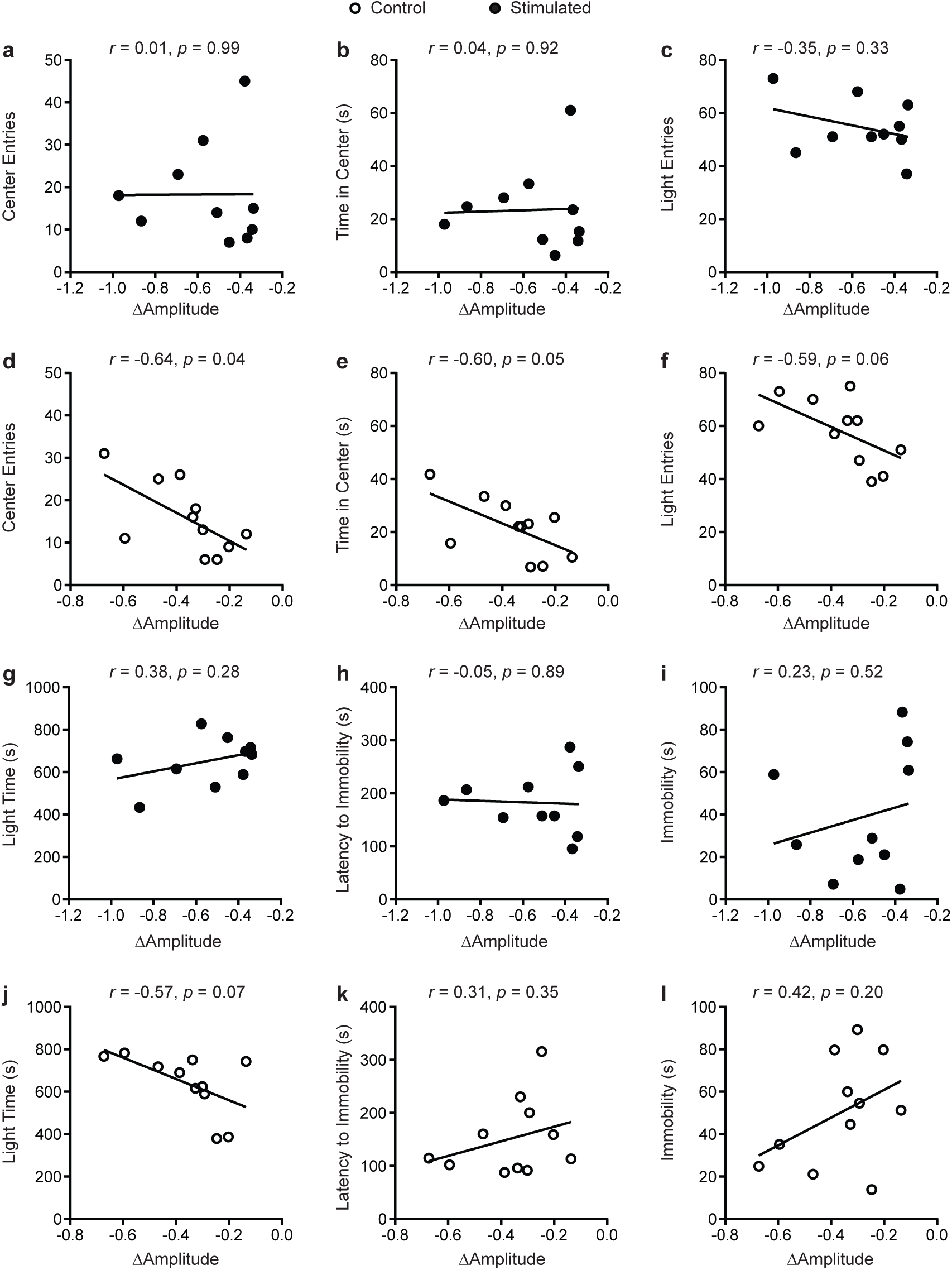
Correlations between change in homecage activity amplitude and behavior in mice that received stimulations or sham stimulations of the SCN at unpredictable times during the dark phase. (a-b, d-e) OF, (c, g, f, j) LD box, and (h-i, k-l) FST behavior. A dampened amplitude of homecage activity rhythms in control mice was associated with decreased anxiety-like behavior in the open field. *n* = 9-11.

One explanation for why homecage activity rhythms are dampened with daily unpredictable dark phase stimulation of the SCN is that our stimulation paradigm disrupts the day/night pattern of rhythms, increasing daily rhythm fragmentation. In addition, we expected that direct stimulation of the SCN during the dark phase would shift rhythms day-to-day (reduce day-to-day stability). To assess daily rhythm fragmentation and day-to-day stability we used a non-parametric circadian rhythm analysis. Similar to the cosinor analysis, the non-parametric analysis revealed that homecage activity amplitude was further decreased in stimulated mice relative to controls (**Fig. 5a**, *t*_(19)_=5.22, *p*<0.0001). Stimulated mice also had a greater increase in homecage activity rhythm fragmentation (**Fig. 5d**, *t*_(19)_=3.55, *p*<0.01) and a greater decrease in day-to-day stability of activity relative to controls (**Fig. 5g**, *t*_(19)_=4.57, *p*<0.001). Consistent with the cosinor analysis, dampened non-parametric activity amplitude was correlated with reduced open arm entries and there was a trend for a correlation with open arm time in the EPM in stimulated mice (**Fig. 5b-c**). Furthermore, increased homecage activity fragmentation (**Fig. 5e-f**) correlated with reduced open arm entries and open arm time in the EPM in stimulated mice. Decreased homecage activity day-to-day stability (**Fig. 5h-i**) was also correlated with reduced open arm entries and there was a trend for a correlation with open arm time in the EPM in stimulated mice. We did not see correlations between the non-parametric circadian rhythm measures and locomotion in a novel environment in stimulated mice or behavior in the EPM in controls (**Supplementary Fig. S6**).

**Figure 5.**
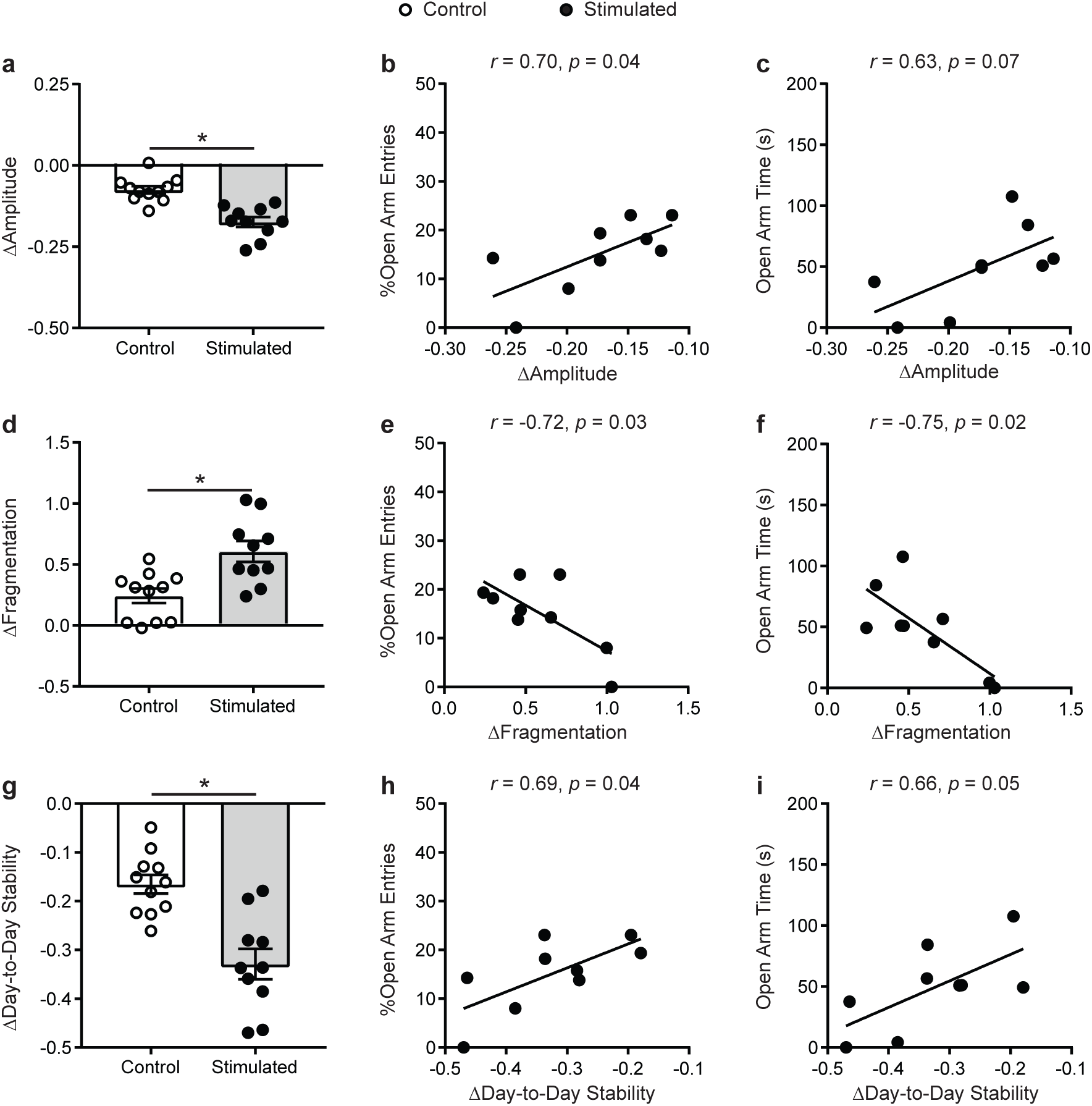
Homecage activity non-parametric circadian parameter changes and correlations with behavior in mice that received stimulations of the SCN at unpredictable times during the dark phase. Change in homecage activity nonparametric circadian parameters were measured relative to baseline in LD. (a) Stimulated mice had a greater reduction in homecage activity amplitude, (d) increase in fragmentation and (g) decrease in day-to-day stability. Correlations between the change in (b-c) homecage activity amplitude, (e-f) fragmentation and (h-i) day-to-day stability with behavior in the EPM in stimulated mice. \**p* < 0.05. *n* = 9-11.

## Discussion

In this study, we used optogenetics to investigate how SCN-mediated chronic advancing and dampening of rhythms affects psychiatric-related behaviors in mice. Of note, this is one of only a handful of studies that have employed optogenetics in the SCN. Stimulation of the SCN at CT21 shortened the period and dampened the amplitude of homecage activity rhythms. Stimulation of the SCN at unpredictable times during the dark phase dampened the amplitude, increased the daily fragmentation and reduced the day-to-day stability of homecage activity rhythms. Neither paradigm significantly altered average daily sleep time, suggesting our SCN stimulation paradigms were primarily affecting homecage activity patterns and not total sleep. In both SCN stimulation paradigms, dampened amplitude of homecage activity rhythms were associated with increased anxiety-like behavior, suggesting that SCN stimulation-induced dampening of rhythms directly increases anxiety-related behavior.

In humans, studies show that circadian rhythms are disrupted in psychiatric disorders^47,48^, but few studies that have examined anxiety specifically. Some studies suggest that environmental circadian disruption, such as by shift work, increases anxiety^15,49,50^. Genetic circadian disturbances, specifically variants in the circadian genes *PER3* and *ARNTL2*, have been linked with anxiety^17,51^. Having a delayed chronotype is also associated with increased prevalence of anxiety disorders^52-54^. One explanation may be that evening chronotypes experience more circadian disruption due to social jetlag, where there are differences between the social and biological clock that results in sleep/wake at inappropriate circadian times. In fact, there is evidence that social jetlag is associated with anxiety symptoms in adolescents^55^. Finally, consistent with our findings in mice, more dampened or fragmented actigraphy rhythms have been found in individuals with anxiety disorders^56,57^. Although there are a limited number of studies on anxiety specifically, work thus far suggests that circadian rhythm disruptions may play a causal role in anxiety disorders in humans.

Both genetic and environmental disruption of circadian rhythms can increase anxiety-related behavior in rodents, but the mechanism is unclear^32,34,58,59^. Our findings suggest that SCN-driven dampening of the amplitude of rhythms increases anxiety-like behavior. Consistent with our findings, we previously found that in stressed mice, more dampened SCN PER2::LUC rhythms were directly associated with increased anxiety-like behavior, suggesting a role for the amplitude of SCN rhythms in regulating anxiety^38^. To determine if disrupting SCN molecular rhythms can cause behavioral changes, Landgraf and colleagues knocked down the core circadian gene *Bmal1* in the SCN and found that it dampened the amplitude of SCN PER2::LUC rhythms and increased anxiety-like behavior in mice^28^. We expect that our optogenetic manipulations of SCN neuronal activity will disrupt SCN gene expression since others have shown that pharmacological disruption of SCN neural activity can dampen SCN gene expression rhythms^60,61^. In future studies, we will determine how SCN optogenetic stimulation affects SCN clock gene expression and if SCN gene expression changes are associated with psychiatric-related behaviors.

Unexpectedly, we did not observe correlations between changes in tau and psychiatric-related behaviors in the mice that received SCN stimulation at CT21. Previous studies showed that exposing animals to an advancing LD schedule for weeks increased depressive and/or anxiety-like behavior^32,33^. We may have seen associations between tau and behavioral measures with a longer stimulation paradigm. However, lengthening our stimulation paradigm would be labor intensive since the animals would be free-running and stimulated by the experimenter at the appropriate circadian time of each mouse. As technology is advancing, in future studies, a closed-loop wireless SCN optogenetic stimulation approach would allow for longer duration paradigms. It is also possible that a longer, but not a shorter tau is associated with anxiety and depressive-like behavior. In humans, an evening chronotype is more consistently associated with adverse mental health outcomes^52,53^ and thus we are interested in determining how SCN-mediated delaying of rhythms would affect behavior.

It is interesting that dampened amplitude of rhythms after repeated SCN stimulation was not correlated with increases in all anxiety-like measures and the specific anxiety-like measures were not consistent across both stimulation paradigms. Although the EPM, OF and LD box tests are all approach/avoidance behavior procedures, there is evidence to support that each individual test measures distinct aspects of anxiety-like behavior. Anxiety-like measures from these approach/avoidance tests are often unrelated^62,63^. Thus, these approach/avoidance behaviors may differentially assess types of anxiety-like behavior (i.e. fear of open, brightly lit or elevated spaces) and the two SCN optogenetic stimulation paradigms used here may have differentially affected these measures. An important point is that the designs of the two experiments were very different (e.g. stimulation every 3 days versus daily, housed in DD versus LD, behavior testing during the active versus the inactive phase, different order for behavior testing) and thus it is not entirely unexpected that we did not observe the same correlations across the two experiments. In addition, the combined effect of shortening the period and dampening the amplitude in one experiment versus primarily dampening the amplitude in the other may have led to distinct neuronal outcomes which manifested in increasing a particular anxiety-related measure. In future studies, it will be interesting to try to tease apart how period and amplitude changes together might impact other behaviors compared to a reduction of amplitude or period alone.

Interestingly, unpredictable sham SCN stimulations resulted in correlations between dampened amplitude of homecage activity rhythms with less anxiety-like behavior in the OF. An important point is that the amplitude measure is the change in homecage activity amplitude relative to baseline. A small reduction in amplitude is expected in control animals due to handling stress and them being removed from PiezoSleep boxes for sham stimulation. It is known that repeated handling can reduce anxiety-like behavior^64,65^. In control mice, having slightly less robust, more flexible rhythms under times of acute stress may be beneficial. There is evidence to support that having a smaller amplitude of SCN rhythms, intercellular desynchrony of the SCN, leads to a more flexible SCN that can adapt to environmental changes^66^. Another possibility is that an SCN-independent mechanism underlies the relationship between the change in amplitude in homecage activity rhythms and anxiety-like behavior in control animals. Acute handling stress through an SCN-independent mechanism may both reduce anxiety-like behavior and the amplitude of homecage activity rhythms. Stress typically dampens homecage activity rhythms, but the effects on SCN rhythms seem to depend on the type and duration of stress^67-69^. In our hands, more frequent and prolonged stress dampened rhythms in the SCN and increased anxiety-like behavior^38^. Thus, there may be an inverse U-shape relationship between homecage activity amplitude and anxiety-like behavior. Having a decreased amplitude of rhythms (either through an SCN-dependent or independent mechanism) may be beneficial to allow for more flexibility under acute stress which would lead to less anxiety-like behavior. However, having a significant SCN-driven dampening of rhythms may increase anxiety-like behavior perhaps through disrupting rhythms in downstream brain regions.

Contrary to what we hypothesized, with SCN stimulation at CT21, more dampened homecage activity rhythms were associated with less immobility in the FST, suggesting that SCN-mediated dampening of rhythms might lead to less depressive-like behavior. It is possible that disruption of SCN rhythms is antidepressant. Knocking out or mutating circadian genes in mice can disrupt locomotor activity rhythms and decrease depressive-like behavior^70-75^. Furthermore, lesioning the SCN in rodents has antidepressant-like effects^30,31^. However, we did not observe a correlation between change in homecage activity amplitude and latency to immobility with stimulation at CT21. In addition, we did not observe correlations between change in homecage activity amplitude and behavior in the FST with stimulation at unpredictable times during the dark phase. Thus, it is also possible that SCN-mediated dampening of rhythms may not play a strong role in regulating depressive-like behavior. In fact, recent work shows that mice exposed to an ultradian LD cycle display increased depressive-like behavior only when ipRGCs that project to brain regions outside of the SCN were left intact, indicating that the SCN does not play a major role in regulating depressive-like behavior induced by light^29^. Moreover, a limitation of this study was that we only used one assay, FST, as a measure of depressive-like behavior. We chose the FST since it is a quick, widely used test, but we recognize that the interpretation and reliability of the FST have come into question^76^; Thus, in future studies it will be necessary to include other depressive-like measures (e.g. sucrose preference or learned helplessness) in separate cohorts of mice.

Ultimately, our findings suggest that SCN-mediated dampening of rhythms can increase measures of anxiety-like behavior in mice. This work is an important step into understanding how the SCN regulates psychiatric-related behaviors. The SCN consists of a diverse population of neurons that receive different inputs, express different neuropeptides, and have distinct neural projections. In future studies it will be important to tease apart the cell-type and circuitry-specific mechanisms underlying the effects of SCN-mediated dampening of rhythms on increased anxiety-like behavior. A more comprehensive understanding of the role of circadian rhythms and the SCN in regulating anxiety may lead to novel chronotherapeutics to treat anxiety disorders.

## Acknowledgements

We thank Drs. Kevin Donohue and Bruce O’Hara for their help with the PiezoSleep software. We thank Dr. Yanhua Huang for sharing equipment and Christine Heisler for helping with our pilot study. This work was funded by grants awarded to Dr. McClung (MH106460, MH115241, MH111601, Brain & Behavior Research Foundation (BBRF) and IMHRO) and Dr. Vadnie (NARSAD Young Investigator Grant from the BBRF). Dr. Vadnie was supported by a NHLBI T32 (HL082610; PI: Dr. Buysse).

