## Supplementary Material for "The Suprachiasmatic Nucleus Regulates Anxiety-Like Behavior in Mice"

### Supplementary Materials and Methods

#### Acute SCN stimulation for c-Fos analysis

Mice were maintained on a 12:12 LD schedule for 1 wk. Mice were then placed in DD for 5-7 days. Stimulation times were determined by finding three consecutive onsets and using linear regression to predict the onset for the stimulation day (day 6 or day 8). Mice were either stimulated or sham-stimulated at CT21 in the dark using a dim red headlamp. Mice were returned to their homecage and perfused 1 hr later under dim red light.

#### Immunofluorescence

Mice were deeply anesthetized with pentobarbital (200 mg/kg, *i.p.*) for c-Fos or ketamine (100 mg/kg, *i.p.*) and xylazine (10 mg/kg, *i.p.*) for VGAT labeling. Mice were perfused with ice-cold 1X PBS followed by 4% paraformaldehyde for 10 min. Brains were removed and postfixed overnight in 4% paraformaldehyde. Brains were cryoprotected in 30% sucrose and cut into 40 μm coronal sections. Floating sections were rinsed in 1X PBS and blocked (5% normal donkey serum, 0.2% Triton-X in 1X PBS) for 1 hr. Sections were incubated with rabbit anti-c-Fos antibody (1:5000, ABE457, EMD Millipore, Burlington, MA) for 48 hrs or goat anti-VGAT (D-18) antibody (1:400, sc-49574, Santa Cruz Biotechnology, Dallas, TX) overnight at 4°C on a shaker. Sections were washed in 1X PBS and incubated with donkey anti-rabbit Alexa 555 (1:500, A31572, Invitrogen,

Carlsbad, CA) or donkey anti-goat Alexa 555 (1:1000, A21432, Invitrogen) for 2 hrs at room temperature. Sections were rinsed and mounted on slides.

Slices immunostained for c-Fos were imaged at 10x with a confocal microscope (FV1200 IX83, FluoView, Olympus, Center Valley, PA). Images were obtained using FV10ASW 4.2 software with a 1.5 zoom. A 30 µm z-stack (with 5 µm steps) was obtained for 3-6 sections containing the SCN per sample (5-6 mice/group). ImageJ was used to count the number of c-Fos-positive cells. Counting was carried out by an investigator blinded to the experimental groups. Sections immunostained for VGAT were imaged at 4x magnification on a fluorescent microscope (Olympus).

**Behavioral Assays**

For all assays, equipment was cleaned and dried in between animals. For behavior testing, we aimed to include ~10 mice per experimental group

**Locomotor activity**

Mice were placed in clear boxes (field dimensions: 9.5” x 18.0”) equipped with photobeams to measure horizontal distance traveled for 60 min (Kinder Scientific Smart Cage Rack System, Poway, CA).

**Open field (OF)**

Mice were placed in the corner of a large plastic arena (52 x 52 x 25 cm) and were allowed to explore for 10 min. The center was a 24 cm x 24 cm square in the middle of the box. Behavior was video recorded and center entries and center time was scored by a blinded, trained observer.

**Elevated plus maze (EPM)**

The EPM consisted of two open arms perpendicular to two closed arms (arms: 30 x 5 cm). Mice explored the maze for 10 min. Behavior was video recorded. Time in the open arms and arm entries were manually scored by a blinded, trained observer.

**Light/dark box (LD box)**

Clear boxes (Kinder Scientific Smart Cage Rack System) were divided into two equally sized chambers, a black opaque chamber kept in the dark and a brightly lit chamber (~880 lux). Mice were first placed in the dark chamber for 2 min and then a door was opened between the two sides. Mice were allowed to explore both chambers for 20 min. Photobeams were used to measure the number of entries into and time spent in the light side.

**Forced swim test (FST)**

Mice were placed into a glass beaker of water (25-26° C; 18 cm depth) and behavior was video recorded for 6 min. Time spent struggling during the last 4 min of the test and latency to immobility were scored by a blinded, trained observer. Struggling was defined as any movement that was not for the sole purpose of keeping the mouse afloat. Immobility time was calculated by subtracting the time spent struggling from the total time (4 min).

### Supplementary Figures

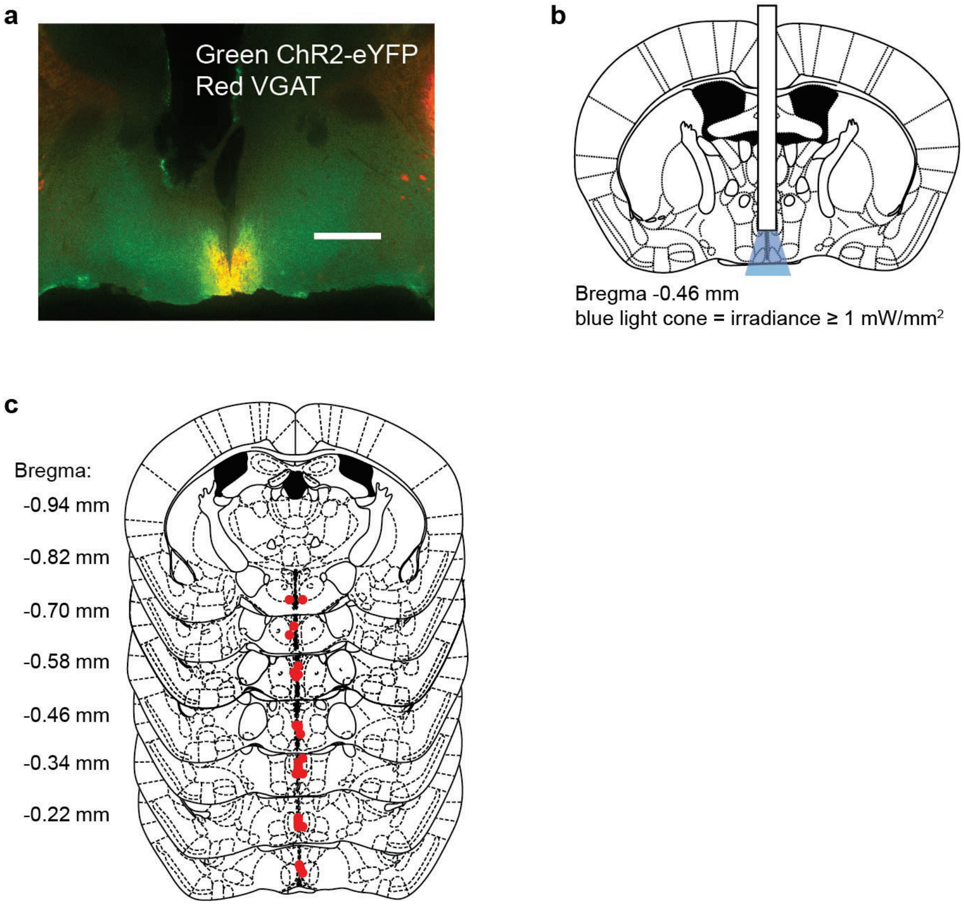


**Supplementary Figure S1**. **Localization of ChR2 in the SCN, blue light spread and placements for mice stimulated at CT21.** (a) Image showing the localization of VGAT and ChR2-eYFP in a coronal brain slice containing the SCN from a *Vgat*;ChR2 mouse implanted with an optic fiber. Scale bar = 500 μm. (b) Illustration of a coronal brain slice containing the SCN showing the ML and DV spread of blue light with an irradiance ≥ 1 mW/mm² emitting from the fiber tip which is sufficient to activate ChR2 (Paxinos and Franklin, 2001). (c) Red points on the coronal brain slices indicate the fiber placements in the SCN of mice that received SCN optogenetic stimulation at CT21.

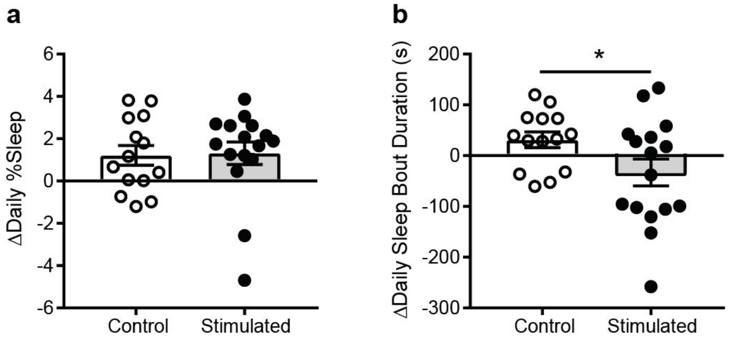


**Supplementary Figure S2. Chronic SCN optogenetic stimulation at CT21 decreased sleep bout duration, but not total sleep time relative to controls.** (a) Change in sleep parameters were measured relative to baseline in DD. Control and stimulated mice exhibited similar increases in average daily % time spent sleeping. (b) Stimulated mice showed a decrease in the change in average daily sleep bout duration relative to control mice (*$p < 0.05$). $n$ = 14-16.

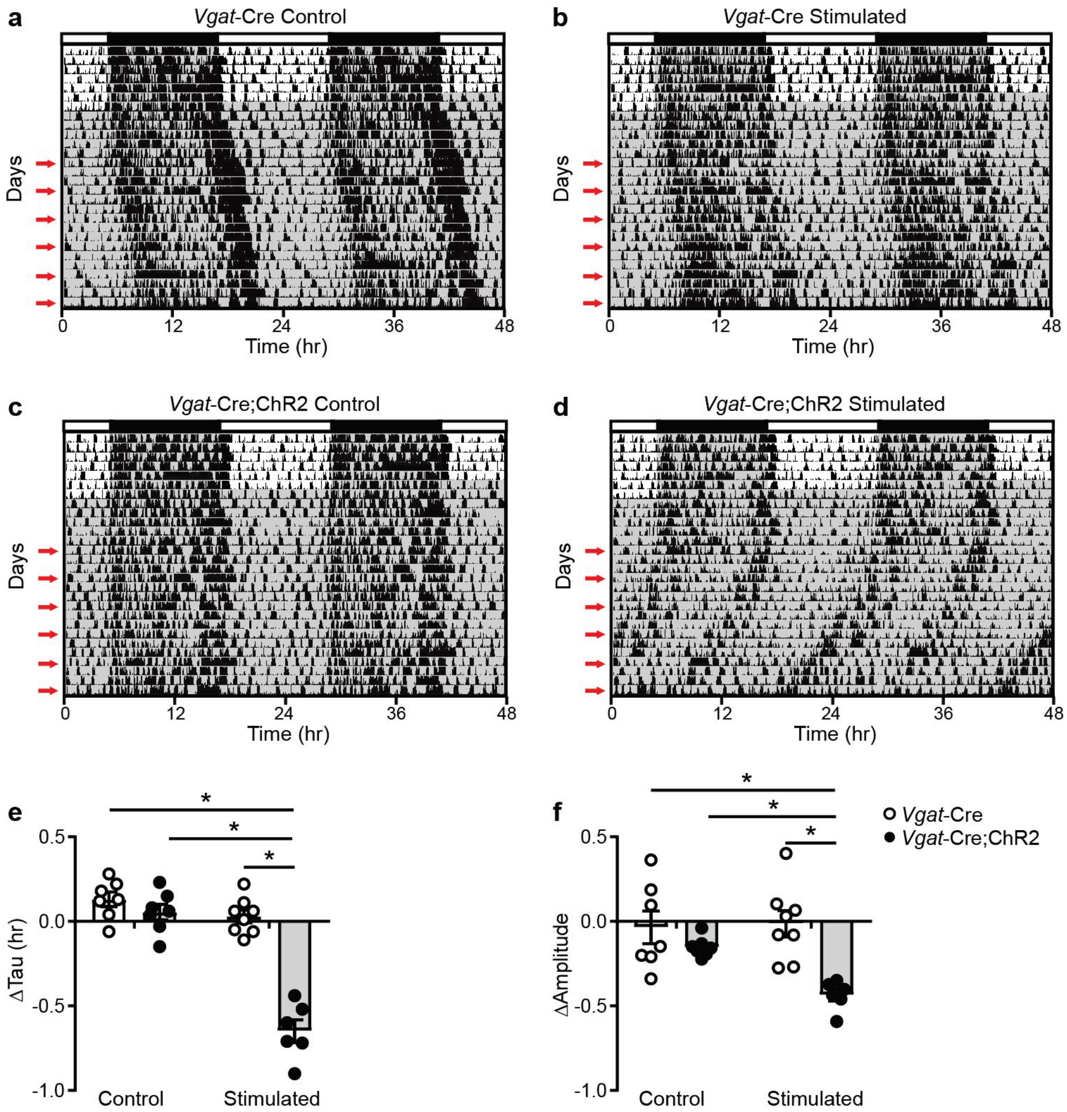


**Supplementary Figure S3. Optogenetic stimulation of the SCN at CT21 decreased the period and dampened the amplitude of homecage activity rhythms in *Vgat*-Cre;ChR2 mice relative to controls.** Representative double-plotted actograms of homecage activity of a (a) *Vgat*-Cre control, (b) *Vgat*-Cre stimulated, (c) *Vgat*-Cre;ChR2 control and (d) *Vgat*-Cre;ChR2 stimulated mouse. Gray shading indicates when lights were off and red arrows indicate the days when stimulations or sham stimulations occurred at CT21. (e-f) Change in homecage activity circadian parameters were measured relative to baseline in DD. Only *Vgat*-Cre;ChR2 mice that received blue light pulses onto the SCN showed a significant decrease in the change in homecage activity tau and amplitude relative to controls (*$p < 0.05$ by Tukey's test).

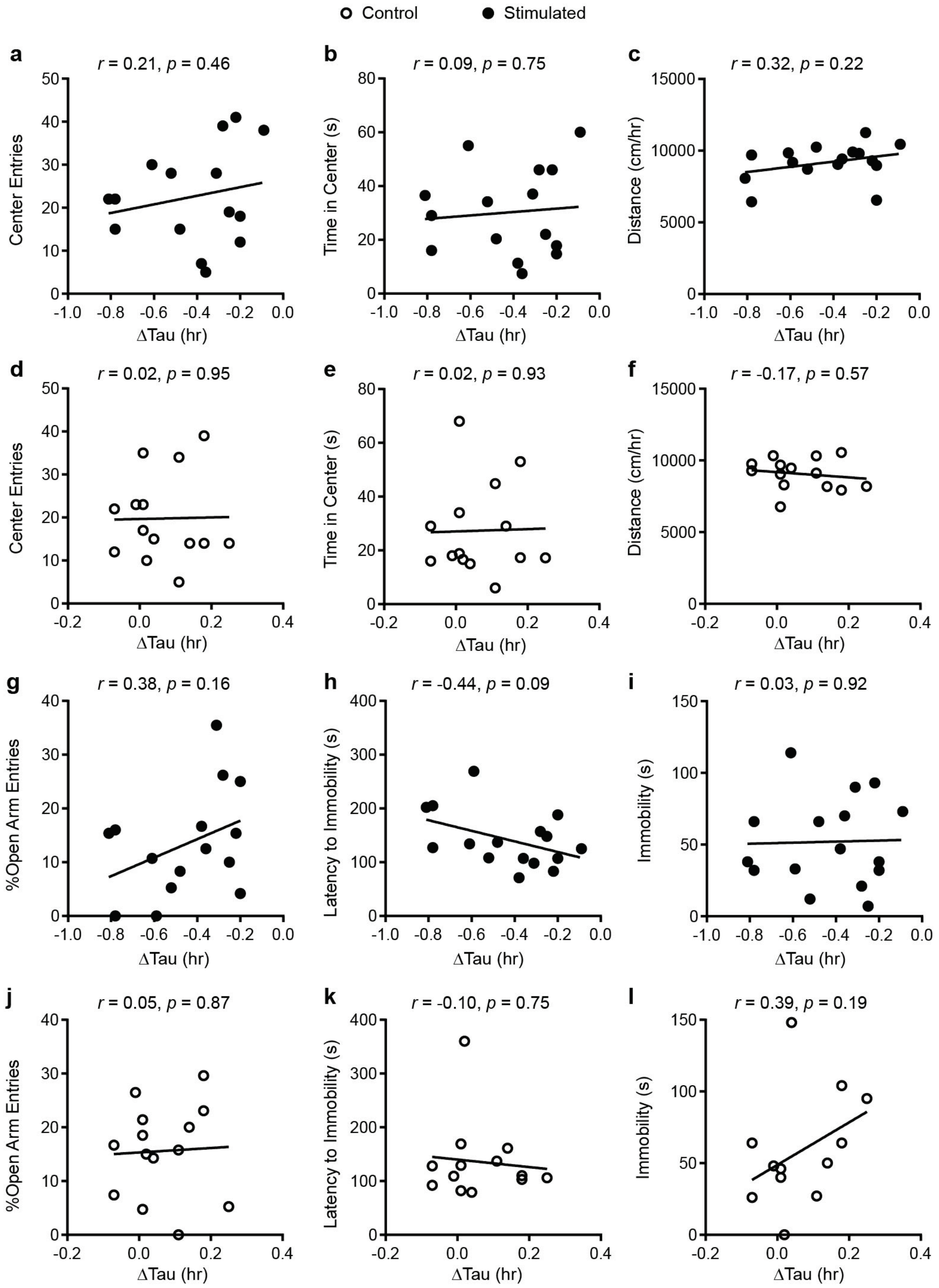


**Supplementary Figure S4. No correlations were observed between the change in homecage activity tau and behavior in mice that received stimulations or sham stimulations of the SCN at CT21.** (a-b, d-e) open field, (e, f) locomotor, (g, j) elevated plus maze, (h-i, k-l) forced swim test. *n* = 13-16.

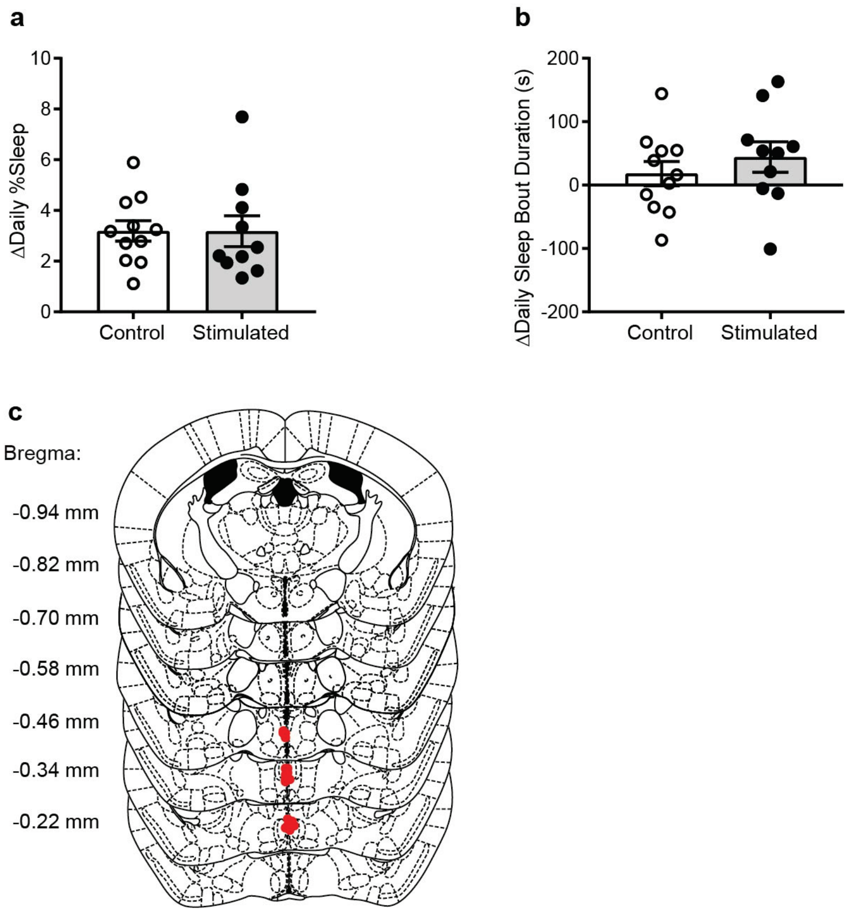


**Supplementary Figure S5. Chronic stimulation of the SCN at unpredictable times during the dark phase did not affect measures of sleep relative to controls.** (a) Change in sleep parameters were measured relative to baseline in LD. Control and stimulated mice exhibited similar increases in average daily % time spent sleeping. (b) Control and stimulated mice also showed similar changes in average daily sleep bout duration. (c) Red points on the coronal brain slices indicate the fiber placements in the unpredictable dark phase SCN optogenetic stimulation experiments. *n* = 10-11.

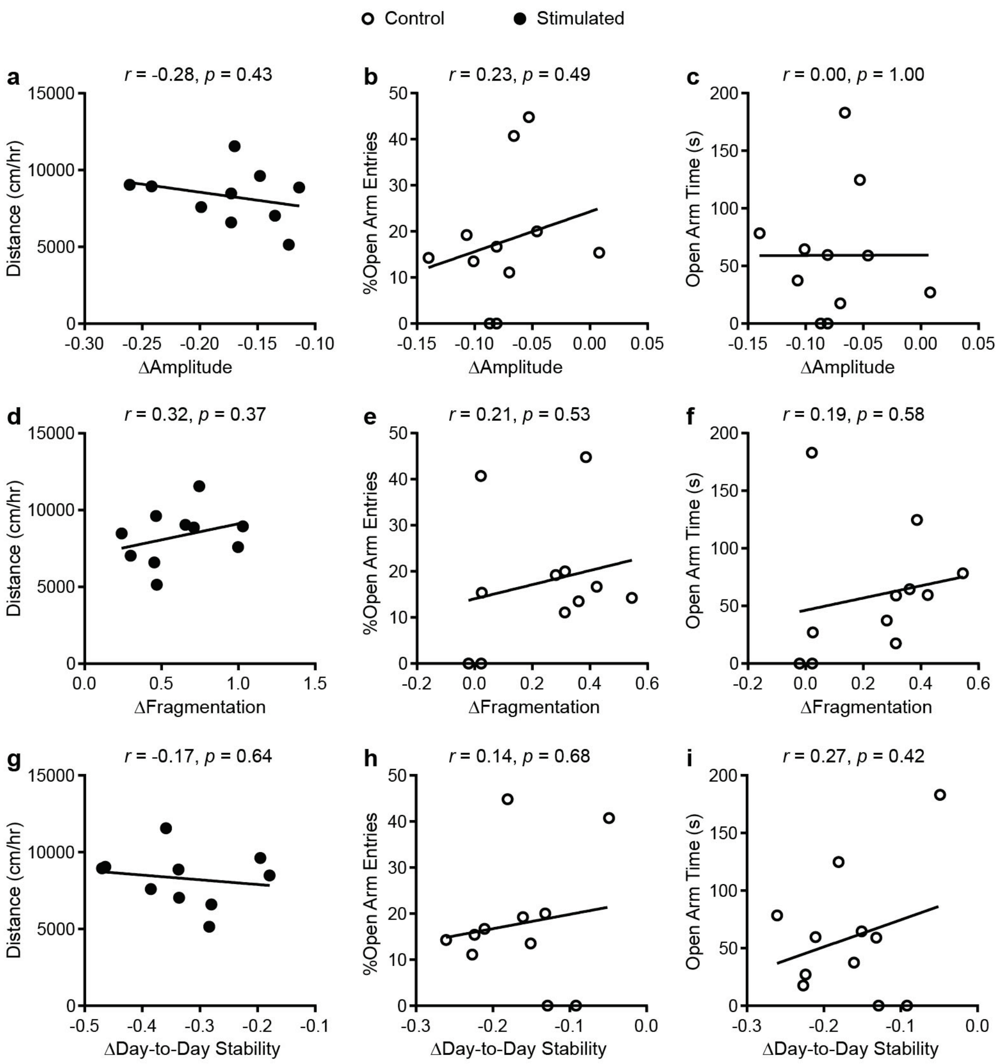


**Supplementary Figure S6. Correlations between non-parametric analysis measures of homecage activity rhythms and behavior.** (a) Correlations between changes in homecage activity amplitude and locomotor activity in a novel environment and (b-c) behavior in the elevated plus maze. (d) Correlations between changes in homecage activity rhythm fragmentation and locomotor activity in a novel environment and (e-f) behavior in the elevated plus maze. (g) Correlations between changes in homecage activity rhythm day-to-day stability and locomotor activity in a novel environment and (h-i) behavior in the elevated plus maze. $n$ = 9-11.
